# Ecology of gene drives: Organism fitness under larval competition determines gene drive performance

**DOI:** 10.64898/2026.11.03.763917

**Authors:** Elliott Hughes, Michael B. Bonsall

## Abstract

Gene drives are a promising technology to control disease vectors, such as mosquitoes. However, as these interventions aim to modify and/or control the population of disease vectors, a full understanding of how gene drives interact with the ecology of the target species is essential. In this study, we explore how larval competition determines gene drive performance. We first extend an agent-based model of the larval behaviour of *Aedes aegypti* mosquitoes to obtain detailed predictions for gene drive designs under realistic larval competition. Using Bayesian optimisation techniques, we explore the performance of different gene drive designs and find the Pareto frontier of optimal underdominant drives with different release thresholds. To understand what determines the structure of the Pareto frontier, we conduct both local and global sensitivity analyses to assess the role different parameters play in determining gene drive performance. We find that ecological parameters, which determine the behaviour of the organism, dominate over genetic parameters for much of the parameter space, although genetic parameters play an important role at the frontier. Finally, we develop stochastic and deterministic surrogate models to enable efficient comparison and simulation without imposing the computational costs of agent-based approaches.

## 1 Introduction

Dengue fever is a widespread arboviral disease transmitted by the bite of infected mosquitoes, with approximately 50–100 million symptomatic cases annually [4, 30]. The cost of these infections is substantial, exceeding $8.9 billion annually (2013 US dollars) [29], with these costs distributed almost evenly between direct medical and productivity costs [16, 28]. The main vector of Dengue fever is *Aedes aegypti*, although *Aedes albopictus* is also an important vector in some settings [5].

As Dengue fever is primarily spread by two mosquito species, and currently available vaccines are projected to only moderately reduce hospitalizations and deaths [32], targeted control of these vector species (especially *Aedes aegypti*) is desirable to interrupt Dengue transmission [1, 37]. A variety of strategies have been proposed, including the introduction of *Wolbachia* parasites and the development of gene drives [1, 15, 32, 37]. Gene drives are selfish genetic elements which spread at a super-Mendelian rate. Several different gene drive architectures have been suggested for the control of *Aedes aegypti* [1, 19, 25, 33]. However, the performance of gene drives is likely to be influenced by the ecology of the target species and in particular by the density-dependent effects which occur during the larval stage of mosquito development [14, 35].

In this study, we explore how competition during the larval phase of the mosquito lifecycle determines gene drive performance. We extend an agent-based model of larval behaviour (first published in [36]) to incorporate a simple but flexible model of a gene drive. Unlike previous work that imposes a generic ‘fitness cost’, we specify the gene drive as imposing penalties on each of the parameters which determine organism behaviour. We allow for varying parameters across individuals which are homozygous and heterozygous for the gene drive allele and incorporate a ‘conversion rate’ to model homing drive architectures. We use this model to design threshold-dependent drive architectures, and use Bayesian optimisation to find the Pareto frontier for efficient gene drive designs (that is, the set of drive designs which most efficiently trade-off the probability of fixation and the probability of extinction at different release thresholds). To understand how different parameters affect gene drive performance, we conduct both local and global sensitivity analyses of the agentbased model. Finally, to enable rapid simulation of this model we utilise machine learning techniques to construct accurate and computationally efficient surrogate models for a gene drive design. These surrogate models could be used to quickly evaluate behaviour under different initial conditions (e.g. with varying initial drive frequencies) or as building blocks for future models that incorporate spatial structure.

### Gene Drives for *Aedes aegypti* Control

To control *Aedes aegypti*, a variety of gene drive architectures have been proposed (including homingbased drives, underdominant drives, and meiotic population suppression drives) [19, 25]. These gene drive architectures can be divided into two broad categories: population suppression drives, which aim to reduce the population size of the target species, and population modification drives, which aim to modify the population of the target species (e.g. by making them less competent vectors for disease transmission) [25]. These gene drive categories can themselves be divided into low threshold drives, which are designed to be ‘fire and forget’ interventions that spread from very low initial frequencies, and high threshold drives, which are designed to be more controllable and will only persist in the population if the initial frequency of the gene drive allele is above a critical threshold [18, 25]. In some cases, gene drive architectures can both suppress and modify the population, especially if the gene drive imposes a significant fitness cost on individuals which carry copies of it [25].

While low-threshold gene drive architectures are more developed for *Aedes aegypti* and offer the highest probability of eliminating the target population, they also carry the highest risk of unintended consequences [19]. Therefore, high-threshold gene drive architectures are actively being developed for *Aedes aegypti* [19]. A common high threshold architecture is the underdominant drive [19]. A variety of mechanisms are available for implementing underdominant drives, including one and two-locus toxin-antidote systems [19]. These underdominant systems are designed such that individuals which are heterozygous for the gene drive allele(s) experience high fitness costs, while individuals which are homozygous for the gene drive allele(s) experience lower fitness costs [9, 35]. This design allows the gene drive to persist if it is introduced above a critical threshold frequency, but prevents the gene drive from remaining in the population if it is introduced below this threshold frequency [3, 6, 31]. If individuals who are homozygous for the gene drive also experience fitness costs then the critical threshold may be high, but this gene drive architecture could be combined with a homing component (that is, a mechanism that overwrites wild-type alleles to bias inheritance in favour of the genetically modified allele) to reduce the release threshold even in the presence of substantial fitness costs [7].

However, the extent to which the gene drive achieves either population modification or suppression (and thus interrupts disease transmission) also depends on interactions between the ecology of the species and the design of the gene drive. In particular, as density-dependence in *Aedes aegypti* is largely mediated by larval competition for resources [15], gene drive efficacy will be partly determined by the fitness costs (if any) that the gene drive imposes on genetically modified larvae [11]. In extreme cases, if a suppression drive reduces the size of the larval population, the consequent reduction in larval competition could result in increased larval survival and little or no reduction in the adult mosquito population [11]. Understanding how these interactions play out is essential for understanding the potential of gene drives to control *Aedes aegypti* and interrupt disease transmission [35]. Consequently, while the development of gene drives for *Aedes aegypti* is progressing rapidly, a detailed understanding of how these gene drives would interact with the ecology of the target species is necessary to understand performance in real-world settings [6, 11, 18, 26].

### Modelling the Interaction between Gene Drives and Larval Competition

Mathematical modelling of the interaction between gene drives and larval competition in *Aedes aegypti* has typically focused on prescribing density dependence by assuming a functional form for the larval survival rate that depends on the local population density. For example, MGDrivE 2.0, a popular software package for simulating mosquito gene drive dynamics, simulates the larval population using an agent-based framework, where individuals transit through a series of discrete life-stages, with transition probabilities dependent on a series of parameterised probability distributions. Density dependence in the larval states is represented by a logistic function, although the package is designed so that more complex functional forms could be used if desired [34]. Similar approaches have been used in ODE-based models, as in the work of Xu and Bonsall [35], and in Markov chain based approaches, as in the model of Otero et al. [24].

Alternative approaches, such as the Skeeter Buster package [22], include emergent density dependence. However, this density dependence occurs at the level of a developing cohort and thus does not fully incorporate interactions between density-dependence and genetics (as genetically modified individuals may perform differently when competing for resources). Alternatively, Hancock et al. [15] introduced density dependence by assuming larval development times depended on density (but that larval mortality per day was constant), so that larvae experiencing intense competition spent more time at risk than those experiencing less competition for resources [15]. Larval density dependence thus emerged from the properties of the system, but the assumption of constant mortality risk may not be realistic under varying food resource availability in the wild.

Beyond *Aedes aegypti*, a variety of approaches have been used to model the interaction of gene drives and density dependence. For example, Depinay et al. [10] constructed an agent-based model for larval competition in *Anopheles gambiae* mosquitoes (see also [23] for another example). Abstract models which are intended to represent general properties of underdominance drives without describing a particular species include work by Champer et al. [6], together with the seminal papers of Barton and Turelli [3] and Tanaka et al. [31].

As prior approaches have relied either on phenomenological models of density dependence or on cohort scale descriptions, neither are fully suitable for modelling the interactions between gene drives and larval competition. However, recent work [36] has developed detailed models of larval competition in *Aedes aegypti* and emergent density dependence. In this work, we build upon these recent developments to study how this emergent density dependence interacts with gene drive behaviour, and thus develop ecologically grounded insights into gene drive architectures.

## 2 Methodology

In this section, we outline the methodology we use to explore how larval competition determines gene drive performance. In particular, we first describe the agent-based model of larval competition that we use to obtain detailed predictions for an individual breeding site (subsection 2.1). Next, we describe the optimisation approach we use to obtain optimal gene drive parameters (subsection 2.2), and the sensitivity analyses we use to understand how different parameters contribute to gene drive success (subsection 2.3). Finally, we describe how we use data-driven techniques to approximate the agent-based model and create a scaleable alternative model (subsection 2.4).

### 2.1 Agent-Based Model of Larval Competition

In this work, we extend an agent-based model of larval behaviour (first published in [36]) to incorporate a simple model of a gene drive spreading through a population at a single breeding site. The model (implemented in Python, although code for simulations without genetics are also available in R, see Appendix A) simulates the life cycle of *Aedes aegypti*, including larval development, pupation, adult emergence, and breeding.

Each larva is represented as an individual agent, which moves and feeds in a spatial grid that represents the breeding site. Feeding is modelled as a competition for resources, where higher weight larvae are assumed to be larger and have a competitive advantage over lower weight larvae. Successful feeders accumulate reserves until they reach a critical reserve threshold (which then leads to pupation), while unsuccessful feeders lose weight and eventually die. The behaviour of larvae is determined by a number of parameters, which can be divided into environmental parameters (which we fix at values corresponding to typical conditions experienced, see Appendix B for details) and individual-specific parameters (summarised in Table 1). Each individual organism’s parameter values are uniformly distributed (except the parameter *c*, see Table 1 for details). For further details of the base model, see [36].

**Table 1:**
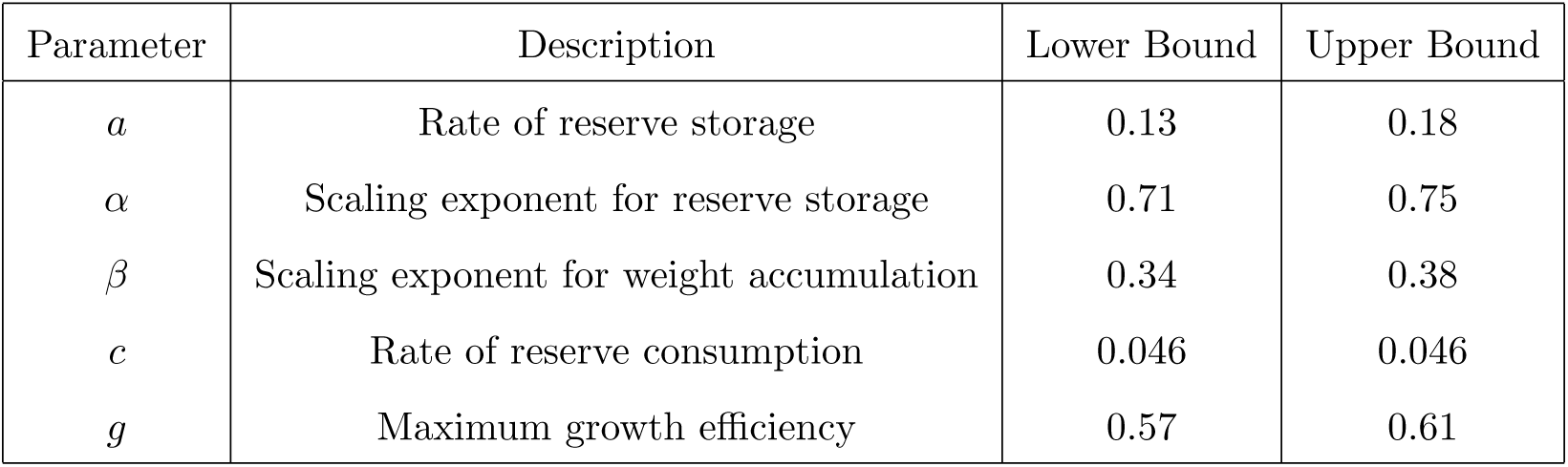
Parameters determining *Aedes aegypti* larval behaviour in the agent-based model (values taken from [36]). For each larvae parameter values are sampled uniformly between the upper and lower bounds, except *c* which is fixed for all parameters.

To extend this model to incorporate a simple description of a single-locus gene drive, we assume that each larva has either W or G alleles at the drive locus, where W is the wild-type allele and G is the drive allele. Organisms with the WG or GG genotype (e.g. heterozygotes and homozygotes for the drive allele) have a fitness cost/benefit that is determined by the impact that carrying these genes have on the individual parameters. While wild type organisms (with the WW genotype) will have parameters distributed uniformly between the upper and lower bounds shown in Table 1, we assume that carrying either one or two copies of the gene drive imposes a penalty on each of the parameters that determine organism behaviour (except *c*, which we keep fixed as in wild type organisms). This is implemented as a multiplicative penalty on both the upper and lower bounds of the distribution for that genotype. The size of this penalty is controlled by a parameter, which we will denote with the base parameter name and a subscript denoting the genotype (e.g. *a*_GG_ for the penalty applied to the distribution of reserve allocation parameter *a*). As an example, if *a*_GG_ = 0.9, then reserve allocation parameters for these GG individuals will be uniformly distributed between 0.9 *×* 0.13 = 0.117 and 0.9 *×* 0.18 = 0.162 (e.g. 0.9 times the upper and lower bounds for wild types).

To propagate the population through time, we simulate the life cycle of the larvae over multiple generations. At the end of each generation, we record the number of larvae that successfully pupate and survive to adulthood. In the absence of a homing component, these adults are divided evenly into males and females, and the next generation of larvae is generated by randomly mating the adults and assigning genotypes to the offspring according to Mendelian inheritance. To incorporate homing drive components, we assume each heterozygote offspring has a probability of converting to a homozygote for the drive allele (we call this parameter the conversion rate). We assume that there is a fixed maximum number of eggs that can be laid at the breeding site, discarding excess eggs randomly if the number of eggs laid exceeds this maximum. This assumption is simple and prevents population collapse due to overproduction of eggs and destructive competition for resources, but assumes that female mosquitoes do not adjust their egg-laying behaviour in response to the density of larvae at the breeding site. In future work more detailed modelling of mosquito egg-laying behaviour could be incorporated, which would enable a more accurate assessment of the strength of the density-dependent competition for resources experienced by larvae at each breeding site. However, given the focus of this study on understanding how larval competition determines gene drive performance, we opt for a simple model of egg-laying behaviour to maintain computational tractability.

In summary, we build upon an agent-based model for mosquito larvae behaviour (first presented in [36]) and extend this model to incorporate a simple description of a one-locus drive. The drive-design parameters that control gene drive performance are the conversion rate and the penalties that the gene drive imposes on individuals with one or two copies of the drive allele. In the sections that follow we will work with a vector of these parameters, which we will define as:

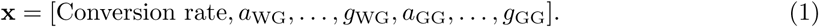

In the following section we will optimise over these parameters to construct threshold-dependent underdominant drives.

### 2.2 Optimisation of Drive Parameters

To understand how these gene drive design parameters determine gene drive performance, we first estimate the Pareto frontier corresponding to the maximally effective high-threshold drive designs for a given critical threshold *τ* (e.g. gene drives designed so that in populations with an initial frequency of the G allele less than *τ* the gene drive becomes extinct, while populations with an initial frequency greater than *τ* the gene drive fixes in the population). To do this, we use multi-objective Bayesian optimisation (see [8] for an introduction to this method).

We consider a population of 100 initial larvae comprised of WW and GG organisms and define the initial frequency of the G allele as *f*_0_ (note that this is also the proportion of GG individuals in the population). For a given set of drive design parameters **x***^∗^*, we then define the frequency of the drive allele in the population at generation *n* as *f_n_*(**x***^∗^, f*_0_). Given a target threshold *τ*, we consider the outcome of a release at a frequency of *τ −* 0.1 and at *τ* + 0.1. The initial frequencies of *τ −* 0.1 and *τ* + 0.1 were chosen to be such that an optimal gene drive design would have a sharp threshold about *τ* (e.g. the gene drive should be such that releases with frequencies close to the *τ* threshold would lead to very different outcomes, with releases just below the threshold leading to gene drive elimination over time and releases just above the threshold leading to gene drive fixation), but so that random variation in breeding outcomes did not significantly impact the results.

As well as choosing the target threshold, we also had to choose the number of generations over which to simulate the model. Choosing five or more generations led to degenerate Pareto frontiers where a large number of gene drive designs always achieved fixation above the threshold and extinction below it. In this case relatively little could be learned about the structure of drive designs or the importance of different parameters in determining behaviour. Conversely, choosing a very small number of generations reduces the influence of the conversion rate parameter, as less breeding cycles occur (in the extreme case, if only one generation is used, no breeding occurs and the conversion rate parameter is meaningless). In our case, preliminary analysis suggested that three generations led to meaningful Pareto frontiers but still allowed the conversion rate parameter to play a role. Thus we define two objective functions *f*_fixation_ and *f*_extinction_ as:

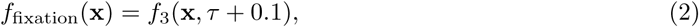

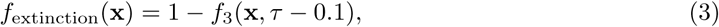

so that *f*_fixation_ corresponds to the frequency of the *G* allele after three generations, while *f*_extinction_ corresponds to the frequency of the *W* allele.

We use the BoTorch [2] library to optimise over these two objectives, using a Gaussian process (GP) surrogate model with a Matern kernel and the q-Noisy Expected Hypervolume Improvement acquisition function. We run the optimisation for 150 iterations, starting from an initial set of 128 points from a Sobol’ sequence in the parameter space (that is, points sampled from a space-filling distribution). To control the noise inherent in stochastic simulations of the agent-based model, we train the GP using the mean and variance of five replicates for each parameter set. To verify function values at the obtained Pareto frontier, we resimulate points identified on the final Pareto frontier a further 20 times to compute a more accurate estimate of the mean and variance of *f*_fixation_ and *f*_extinction_ at these points. To account further for variability, we report Pareto frontiers for five separate Bayesian optimisation runs, extracting the best performing gene drive designs from the combined set of Pareto optimal points. Further implementation details can be found in Appendix C.

### 2.3 Sensitivity Analysis

In addition, we conduct sensitivity analyses to understand how changes to each of the drive design parameters (e.g. *a*_WG_, *a*_GG_, *b*_WG_, etc) affect the the performance of the drive. Firstly we conduct a global sensitivity analysis by calculating the Sobol’ indices for each parameter and objective function. These Sobol’ indices measure the proportion of the variance in the objective function that can be attributed to each parameter, both individually and in combination with other parameters. We calculate the Sobol’ indices (using 20 *×* 2048 = 40960 samples) for each parameter and objective function, which allows us to identify which parameters have the greatest influence on gene drive performance, and how they interact with each other.

These Sobol’ indices provide estimates for the global influence of parameters on outcomes and suggest which parameters are most or least important (e.g. have the largest or smallest Sobol’ indices). However, preliminary analysis suggested that the influence of parameters on outcomes may differ across the parameter space (e.g., the conversion rate may have a low influence on outcomes in regions of the parameter space where the *GG* organisms have very low fitness and do not survive to reproduce). To test the influence of parameters locally to points on the Pareto frontier, we conduct a local sensitivity analysis around the point with the highest average score across both objectives. In particular, we estimate the derivative of each objective function with respect to each parameter (that is, the rate of change of the objective function when the parameter is perturbed by a small amount). We use central finite differences with a perturbation of 10*^−^*^1^, computing the average effect over 100 replicates for each parameter to reduce the impact of random variation, to produce an estimate of the derivative:

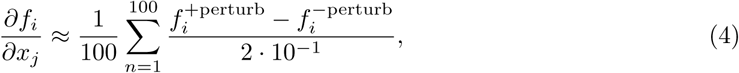

where *∂f_i_/∂x_j_*is the derivative of the objective function *f_i_* with respect to the parameter *x_j_* and 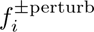 is a realisation of the objective function when *x_j_* is perturbed by *±*10*^−^*^1^, respectively. In the case where one of 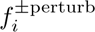 cannot be computed because the perturbed parameter would be outside the biologically relevant bounds (e.g. for parameter sets with a zero conversion rate) standard one-sided finite difference estimates are used instead.

### 2.4 Approximation of the Agent-Based Model

Given the Pareto frontier of optimal gene drive designs identified in subsection 2.2, we use data-driven techniques to approximate the agent-based model. These data-driven approximate models can then be simulated with a single function call, reducing computational costs compared to the multi-step process required to evaluate an agent-based model. We select the gene drive design with the highest average score across both objectives, and train both deterministic and stochastic surrogate models to predict the number of larvae of each genotype that survive to adulthood given an initial population of adults laying eggs into a breeding site. Formally, we define:

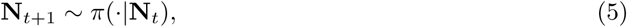

where **N***_t_* = [*N_W W,t_, N_W G,t_, N_GG,t_*] is the number of adults of each genotype at generation *t*. We generate 40000 samples from this distribution by randomly generating different adult populations (which vary in total size and in the number of individuals of each genotype) and recording the resulting number of larvae of each genotype that survive to adulthood. For the deterministic surrogate model, we train a feedforward neural network (which we will denote *f ^NN^*) to predict the mean of this distribution and obtain a mapping:

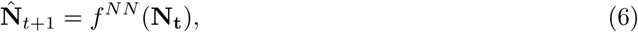

where **N̂***_t+1_* is the predicted number of surviving larvae of each genotype. This mapping can then be used to predict the mean trajectory of the population. However, in order to conduct simulations over more than one generation, we must make a moment-closure assumption that:

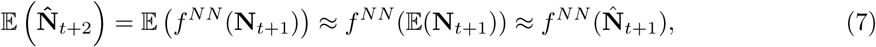

and this assumption is less likely to be true when the variance of **N̂***_t+1_* is high (as in general the mean of a non-linear function of a random variable depends on both the mean and variance of the variable).

For the stochastic surrogate, we train a neural spline flow (NSF) to approximate the observed distribution of outcomes arising from the agent-based model (e.g. that given by Equation 5) with a surrogate distribution *π̃*:

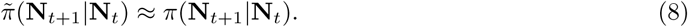

Implementation details for the deterministic and stochastic surrogate models can be found in Appendix D and Appendix E, respectively.

To test how accurately these approximate surrogate models can reproduce the observed dynamics of the agent-based model, we conduct both single-generation and multi-generation simulations of the model to generate ‘test’ datasets that can be used to measure surrogate performance.

For the single-generation tests, we generate a test dataset of 2000 simulations of the agent-based model using the same procedure we used to generate the training dataset. To measure the accuracy of the deterministic surrogate, we then compute the surrogate model’s prediction for each of the 2000 simulations and calculate the root-mean-square error (RMSE):

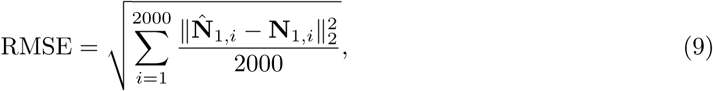

Where *N̂_l,i_* is the model prediction after one generation for the i-th sample in the test dataset and **N**_1*,i*_ is the observed outcome of the agent-based simulation.

To assess the accuracy of the stochastic surrogate model we use the same dataset with an distribution-based testing framework first presented in [20]. In this framework, the tester constructs a function *π̃^−^*^1^ (further details on the construction of this function can be found in Appendix F) such that:

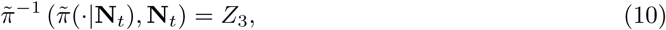

where *Z*_3_ is the three-dimensional standard normal distribution. Applying this transformation to the test dataset yields a set of points which can be compared to a standard normal distribution to assess how well *π̃* approximates *π* [20]. For example, if the mean of the transformed test dataset is non-zero, then this suggests that the surrogate produces biased output, while if the standard deviation of the transformed distribution is not [1, 1, 1] then the surrogate over or under-predicts the variability of outcomes for one or more of the genotypes.

For the multi-generation test, we simulated fifty replicates of the agent-based model for ten generations (beginning with a fixed initial condition of 50 WW and 50 GG organisms), and calculated the mean trajectory. For the deterministic surrogate model, we compared the model prediction to this mean trajectory, while for the stochastic surrogate model we simulated fifty replicates to assess both the mean trajectory of this surrogate model and the variability of both the agent-based model and the stochastic surrogate.

## 3 Results

In the following sections, we present the results of our analysis. We first consider the results of our optimisation of gene drive parameters (subsection 3.1), followed by our sensitivity analyses (subsection 3.2), how these results change if the target threshold is modified (subsection 3.3, and finally the accuracy of our approximation methods (subsection 3.4).

### 3.1 Optimisation of Gene Drive Parameters

Beginning with a fixed target threshold of *τ* = 0.4 (near the threshold for spatial spread identified by [31]) Bayesian optimisation of the gene drive parameters identified a Pareto front of optimal gene drive designs. There was significant variation between different replicates of the Bayesian optimisation process, but the resulting Pareto frontiers were qualitatively similar (see Figure 1).

**Figure 1:**
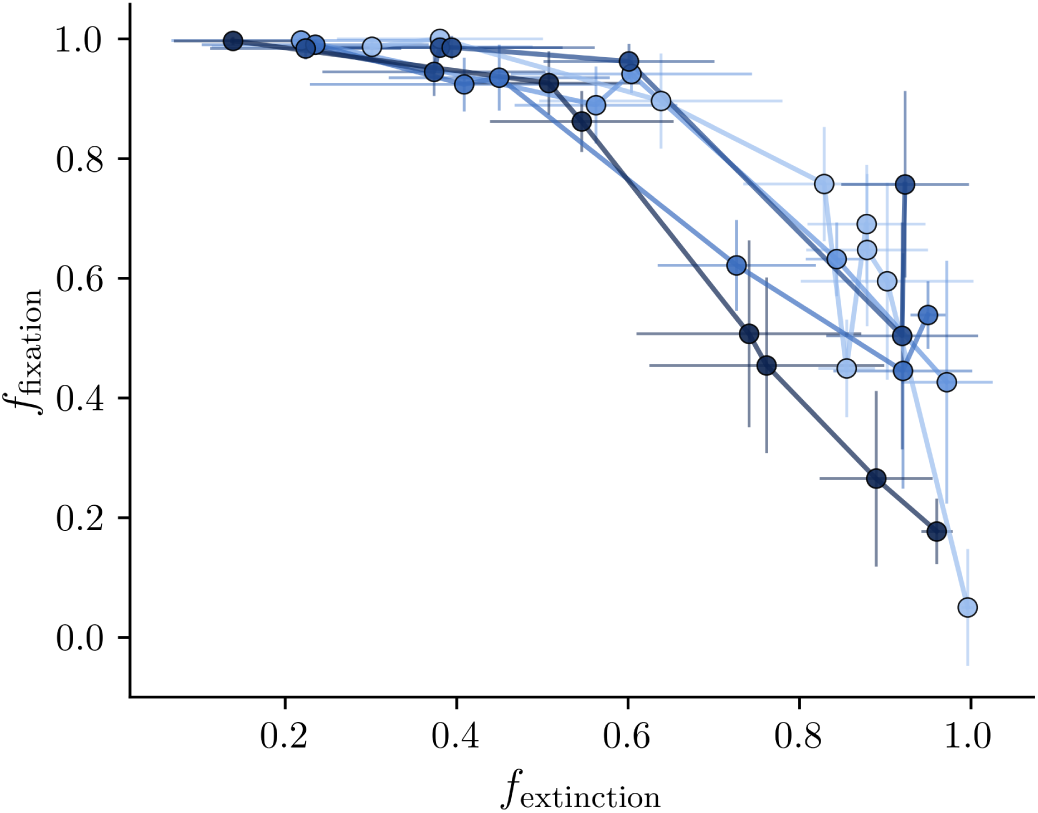
Pareto frontier of optimal gene drive designs identified through Bayesian optimisation (each colour represents a distinct ‘run’ of the optimisation process). Each point represents the mean outcomes of 20 replicates of a set of gene drive parameters, with the x-axis representing the average end frequency of the W allele with a low initial proportion of GG organisms and the y-axis the end frequency of the G allele with a high initial proportion of GG organisms. Thin lines show *±*2 standard deviations around the mean outcomes.

To understand the structure of optimal gene drive designs, we extracted the top five highest performing points on the Pareto frontier and considered the resulting parameter sets. Preliminary analysis suggested that these points could be divided into two clusters, corresponding to lower and higher values of *f*_extinction_. Consequently, in Figure 2 we present the average and individual parameter values within each cluster. We observed significant variation in parameter values within each cluster (particularly for the parameters determining heterozgyote performance and the conversion rate), suggesting that many parameters play only a small role in determining gene drive performance and/or that there are multiple parameter sets which produce similar functional effects. Surprisingly, we observed significant heterogeneity in the conversion rate associated with optimal designs, suggesting that some gene drive designs used a homing component to balance fitness costs, while others had a fitness advantage for individuals with two copies of the G allele. Furthermore, as observed in Figure 1, the noise arising from the stochastic nature of the agent-based model was on the order of 10% of the observed variation in either objective function. This suggests that the impact of small differences in parameter values may be difficult to observe, as the effect of these small changes may be swamped by stochastic variation.

**Figure 2:**
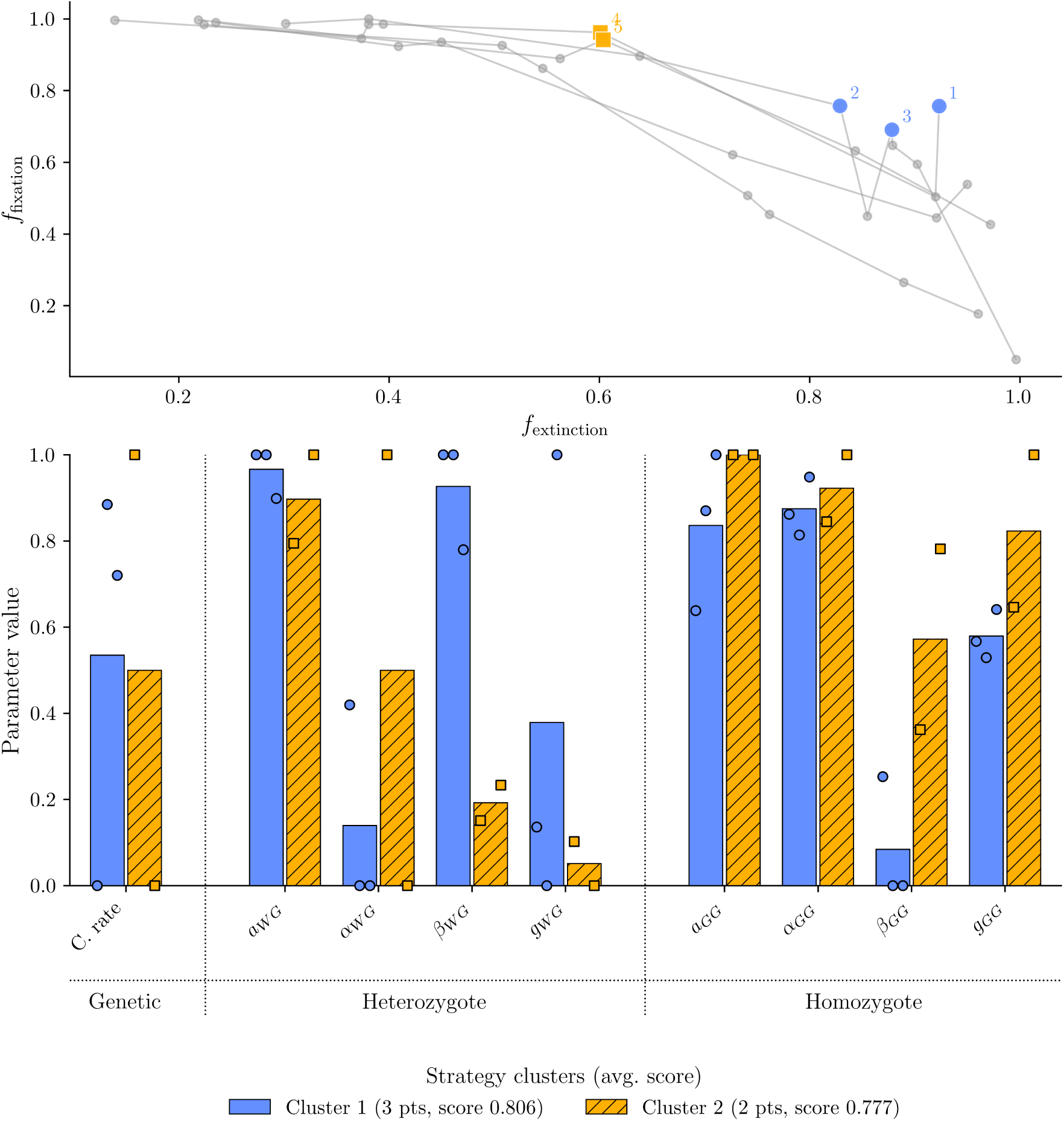
Parameters of the top five optimal gene drive designs identified through Bayesian optimisation. Points are clustered and the mean parameter values are shown for each cluster (bars) with individual parameter values shown as circles (for cluster 1) and squares (for cluster 2).

To understand how these gene drive designs modify larval behaviour to create threshold-dependent dynamics, we began by ignoring the effects of competition and considering the growth trajectories of larvae with access to unlimited food (see Figure 3). All optimal designs featured underdominant dynamics and non-viable heterozygous larvae, with either larvae unable to assimilate resources (leading to immediate starvation), or larvae that allocate almost all resources to increasing total weight and size rather than accumulating reserves (and thus do not achieve pupation within the maximum time allotted by the simulation). In a setting with constrained resources such larvae also likely starve if they do not succeed in obtaining resources, as these larvae will quickly exhaust their limited reserves. Notably, this suggests the significant variation in parameters controlling the behaviour of heterozygotes observed in Figure 2 reflects the multiple distinct ecological ways that larvae can be made non-viable rather than different gene drive designs attempting to induce different behaviour/physiological effects in heterozygotes.

**Figure 3:**
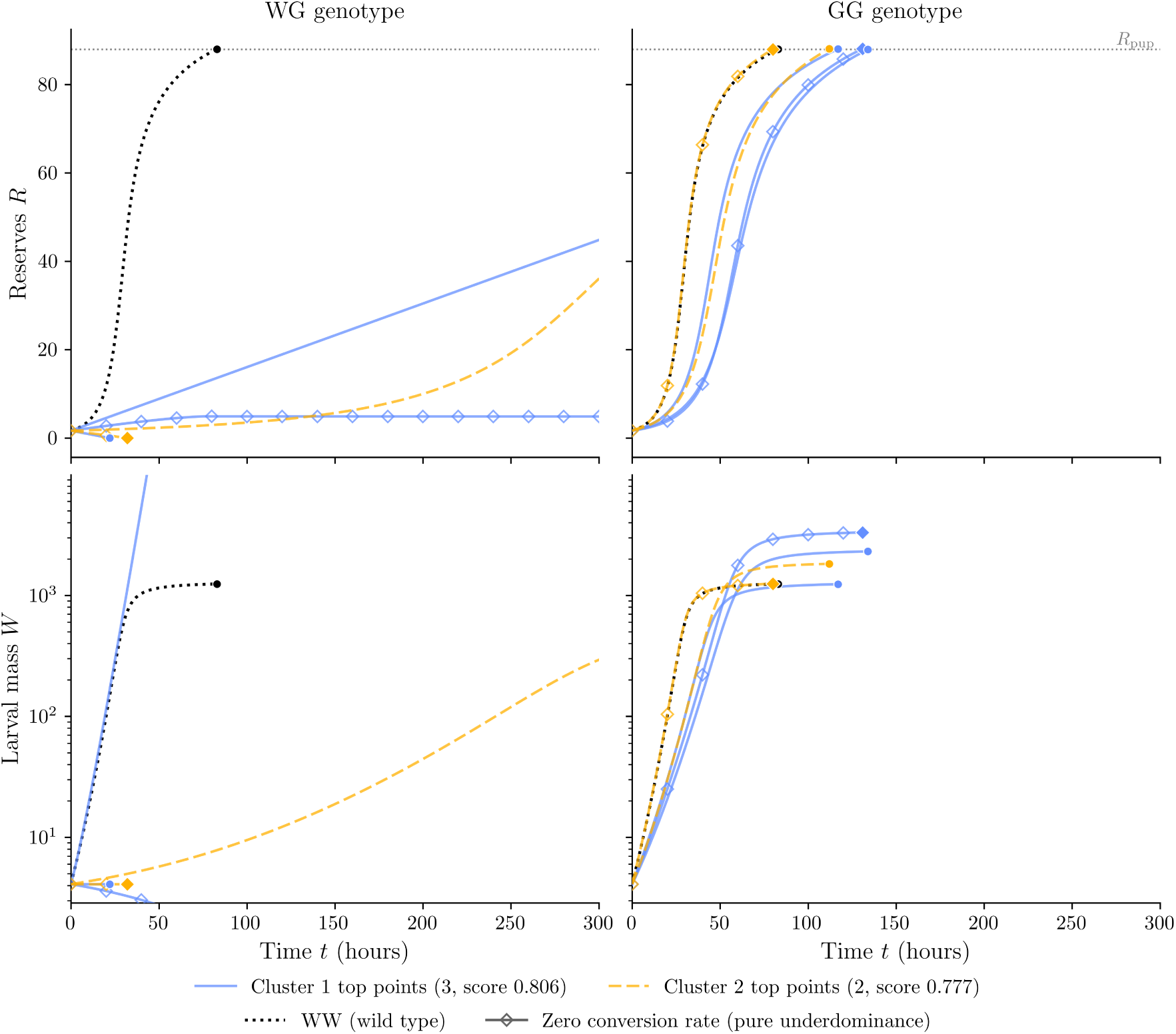
Mass (*W*) and reserves (*R*) trajectories over time for heterozygotic (WG) and homozygotic (GG) larvae, assuming unlimited food access, coloured by cluster membership. Wild-type dynamics (black dotted line) are shown as a comparison, and gene drive desgins corresponding to gene drives with zero conversion rate are highlighted. Trajectories are plotted until starvation *R* = 0, pupation *R* = *R*_pup_, or the end of the cycle, whichever comes first.

The majority of gene drives had low *g*_GG_ and *β*_GG_ values, implying slower growth rates and a higher allocation of resources to increasing total weight rather than reserves. Overall, this design slowed development, but allowed maximum weights to exceed those of wild-type organisms. Thus, when competing with wild-type organisms, this increased allocation to weight and size gain likely gave GG genotypes a competitive advantage during the latter part of the simulation (as in our model higher weight individuals are more likely to succeed in gaining resources, see subsection 2.1). In particular, for the zero conversion rate (e.g. pure underdominance) design, this increased allocation to weight gain appears to have been the main mechanism by which a fitness advantage for the genetically modified organisms was obtained.

Our results indicate that a variety of optimal gene drive designs exist, but that the most successful incorporated strong suppression of heterozygous larvae. Optimal gene drive designs consistently allocated more food to increasing weight than to increasing reserves, delaying pupation in exchange for increased competitive strength during the late phases of development. For most gene drive designs, this was combined with a non-zero conversion rate, but the gene drive with highest overall score purely utilised this increased allocation to weight gain in order to achieve underdominance and a critical release threshold below 50%.

### 3.2 Sensitivity Analysis Results

The analysis above gives insights into the optimal construction of gene drives at the Pareto frontier, but compares sets of parameters rather than measuring the individual impact of each parameter on each objective function. To understand the impact of each parameter on gene drive performance, we sampled a large number of parameter sets (approximately 40,000 in total) and calculated approximate Sobol’ indices for each parameter and objective function. The total Sobol’ indices (see Figure 4 below) show the proportion of the variance in each objective function that can be attributed to each parameter, suggesting how important that parameter is in determining gene drive performance.

**Figure 4:**
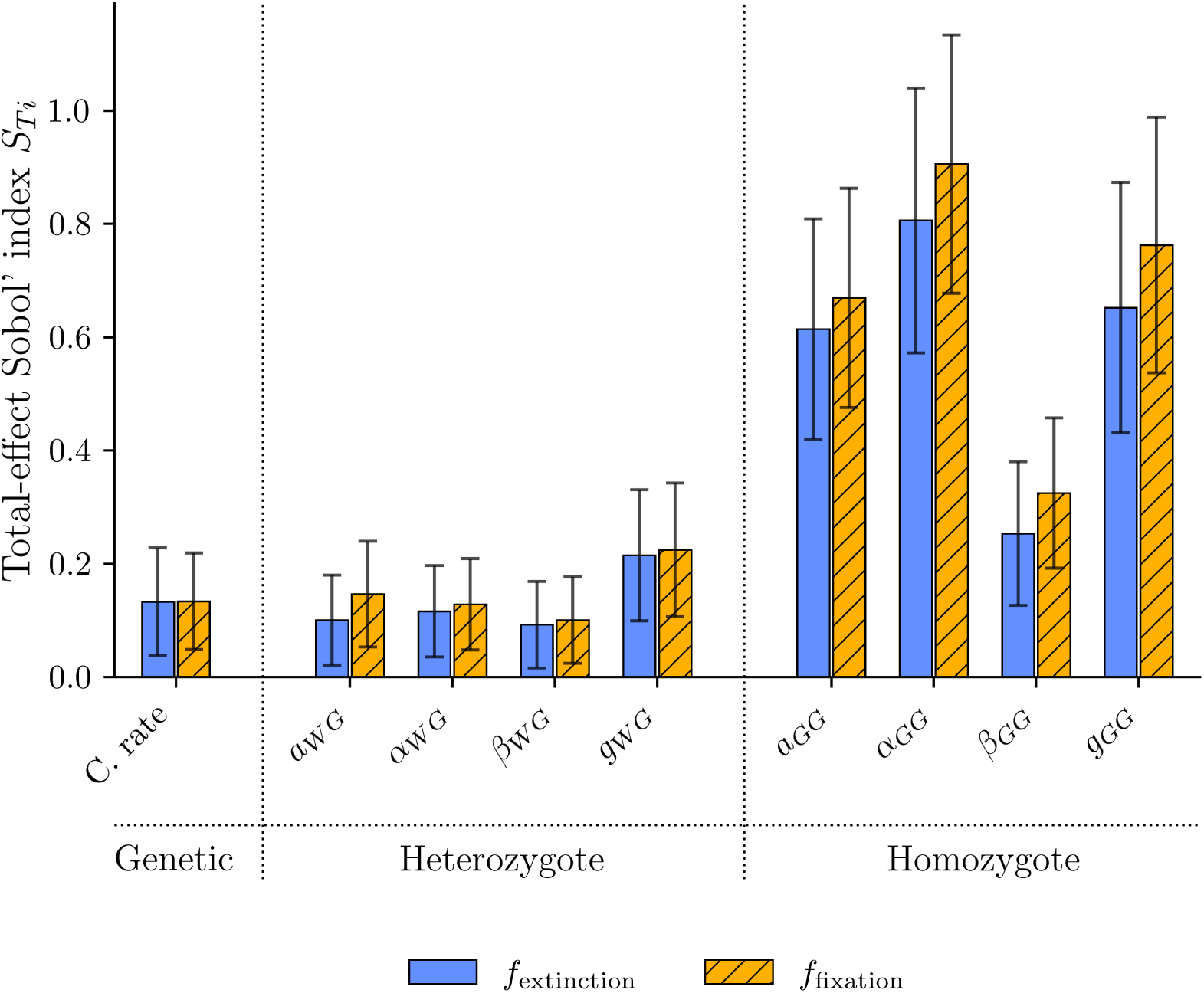
Global sensitivity results for each objective function and each gene drive design parameter. The height of each bar gives the total Sobol’ indices for each parameter and objective function, so that a larger bar suggests that the parameter plays a larger role in determining drive performance. Black whiskers show a bootstrapped *±*2 standard deviation interval around the central estimate, and the parameters are grouped into categories according to their role.

This global sensitivity analysis revealed that the primary determinants of gene drive performance were the penalties imposed on organisms homozygous for the drive allele, suggesting that in many regions of the parameter space the main determinant of gene drive performance was the behaviour of GG organisms. Other parameters, most notably *g*_WG_ also appear to play a secondary role in determining the efficacy of a gene drive. Since there are large regions of the parameter spaces where fitness costs on larvae homozygous for the gene drive are high, these results demonstrate that ecological and behavioural parameters for gene drive individuals are far more likely to determine whether the gene drive is viable, as without viable GG organisms no gene drive design will achieve success. Analysis of the second order Sobol’ indices (which measure the interaction effects between parameters) highlighted the importance of the parameters controlling GG organisms’ behaviour, further supporting this finding (see Appendix G).

Since this global sensitivity analysis considers results over a large range of regimes, many of which are not viable gene drive designs, we also conducted a local sensitivity analysis to understand gene drive behaviour around the gene drive design with the highest average score weighting both objectives equally (see Figure 5).

**Figure 5:**
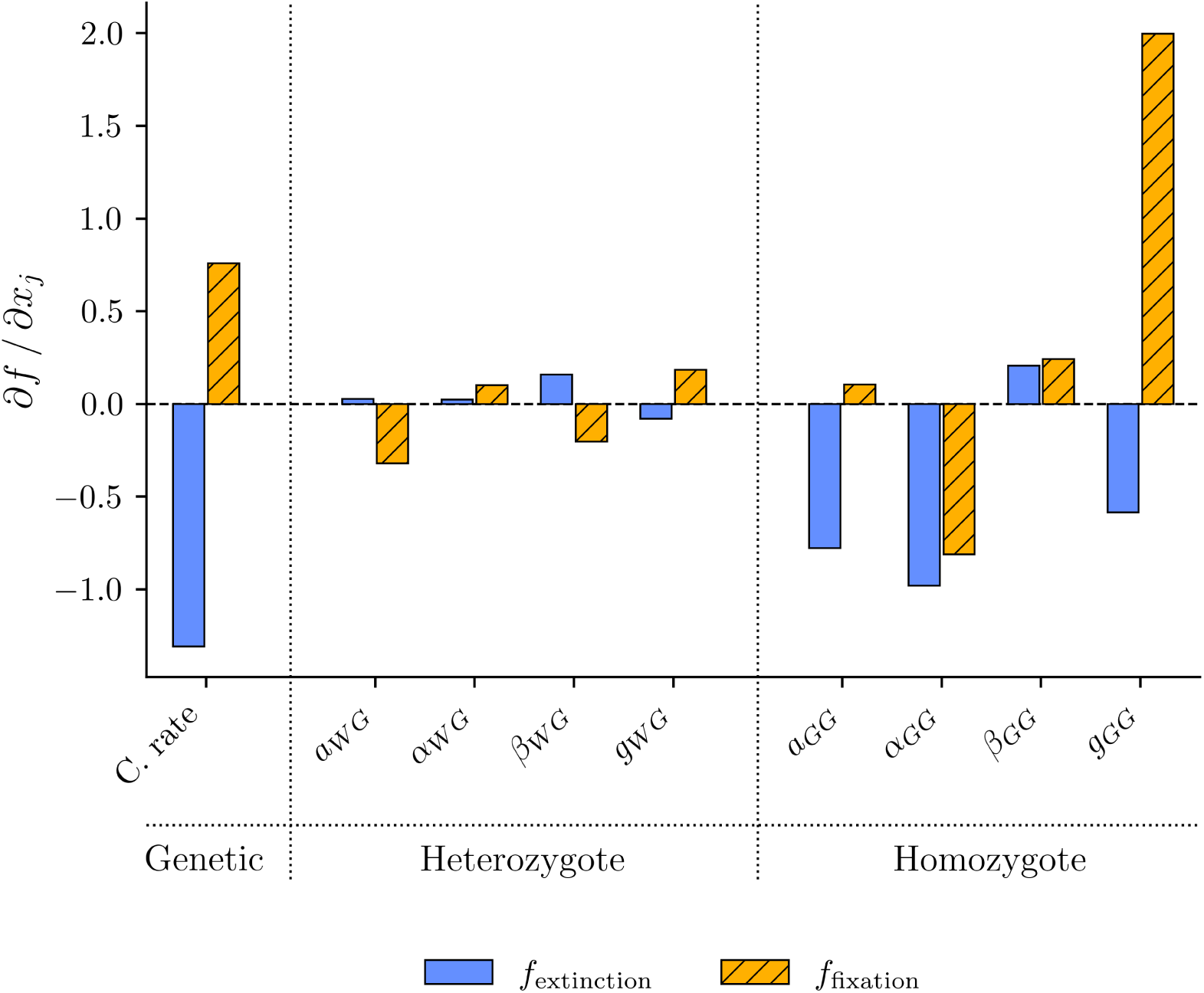
Local sensitivity analysis of gene drive performance around the gene drive design with the highest average score. Each bar represents the average of 100 replicates of a central finite difference estimate of the partial derivative of the objective function with respect to that parameter at the gene drive design.

In this underdominant regime, the primary determinants of gene drive performance were the conversion rate and the maximum growth efficiency for gene drive organisms (*g*_GG_). The conversion rate had a negative effect on gene drive performance (with respect to the extinction objective), significantly reducing the threshold for fixation and thus making it more difficult to find a set of parameters that would lead to gene drive extinction below the release thresholds.

The maximum growth efficiency for homozgyotes for the gene drive also had a significant impact on gene drive performance, with higher values of *g*_GG_ leading to better performance on the fixation objective but worse performance on the extinction objective. This suggests that, at least locally to the point we analysed, altering this parameter also had the effect of moving gene drive designs along the Pareto frontier. This result is unsurprising, as increasing *g*_GG_ will increase fitness for homozygotes of the gene drive.

We also observed non-trivial coefficients for the remaining behavioural parameters associated with GG organisms, and significantly smaller effects for changes to the parameters controlling heterozygotes behaviour. This suggests that for this top-scoring underdominant drive design, heterozgyote fitness is very low, so that small changes in any of the parameters controlling heterozgyote behaviour are insufficient to restore the viability of these larave.

In summary, the results of our global sensitivity analysis reveal that ecological/behavioural parameters determine the effectiveness of gene drives over much of the parameter space we explored (as these parameters determine the fitness costs experienced by homozygotes and heterozgyotes). However, local to the Pareto frontier, where gene drive designs are well optimised to achieve threshold-dependent dynamics, both genetic and ecological parameters played an important role in determining behaviour.

### 3.3 Varying Release Thresholds

The above results consider gene drives with a fixed target release threshold of 0.4. Drives with this release threshold would be expected to spread slowly in a realistic spatial environment [31] which may or may not be a desired property in real world settings. To understand how drives with different release thresholds might behave, we now consider how results differ under varying release thresholds. To understand how parameter sets change when the target release threshold is modified, we consider target critical thresholds ranging from very low (*τ* = 0.2) to very high (*τ* = 0.8). We extracted the three highest scoring points for each release threshold, and show both the individual and mean parameter values for each parameter in Figure 6

**Figure 6:**
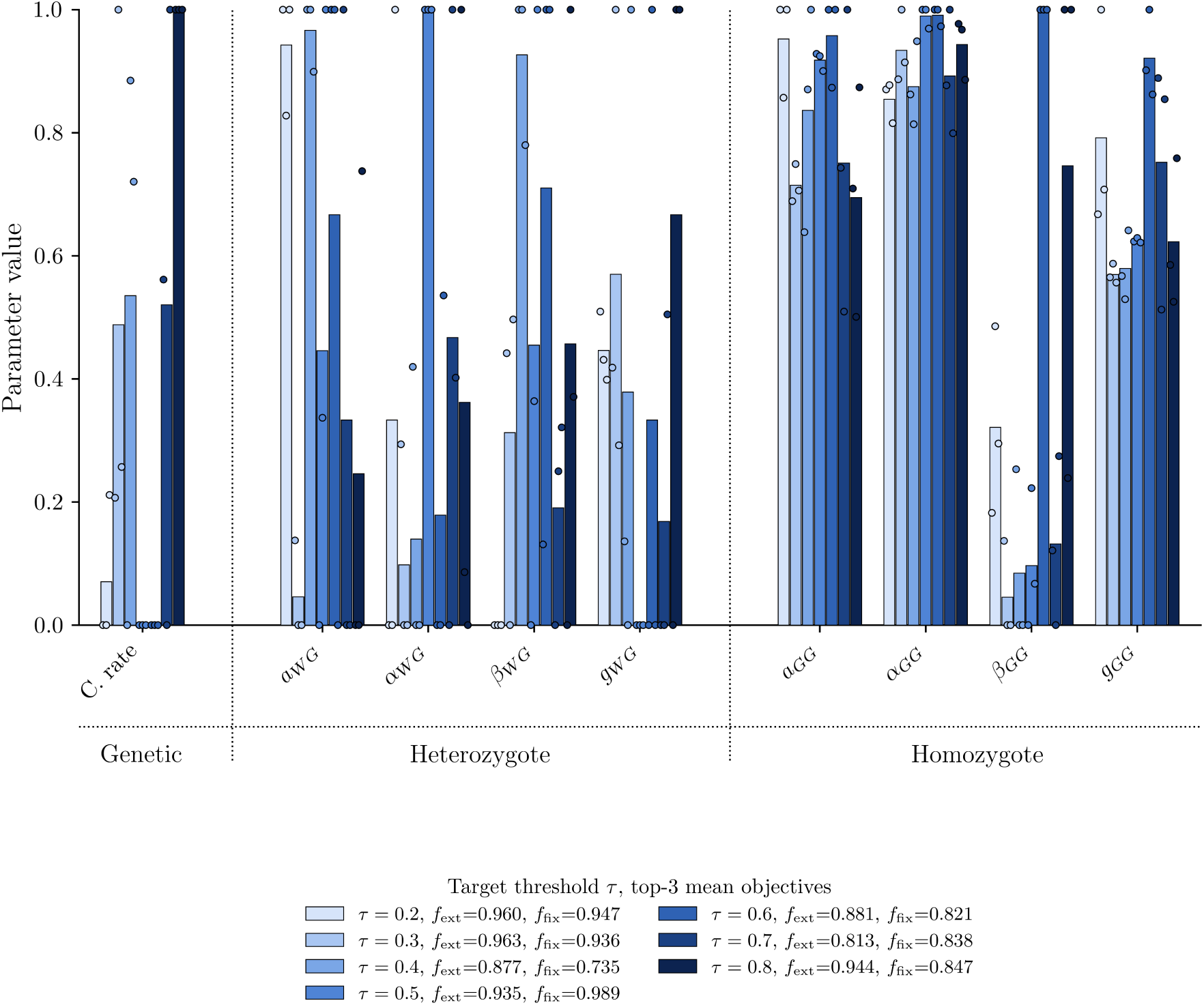
Parameter values for the top three highest scoring drives for each pair of release frequencies (coloured light to dark in order of increasing *τ*). The mean (bar) and the individual parameter values (circles) are shown, highlighting the variability in optimal designs for each value of *τ*. Note that for intermediate values of *τ* the optimal conversion rate is zero, while gene drives designed for very high and very low critical thresholds have non-zero conversion rates.

As with the fixed release threshold we considered above, we see significant variation in parameter values within and between each threshold, suggesting that there are multiple ways to design high-scoring gene drives for each release threshold. However, the conversion rate parameter was uniformly zero for intermediate release thresholds, suggesting that for these cases (where we might expect underdominant dynamics to produce similar results even without a fitness cost for the GG organisms) such purely underdominant designs were the most efficacious.

To understand how the optimal gene drive designs function, we again plotted larval weight and reserves over time if given access to an indefinite amount of food. We find that gene drive designs are very similar to those in the previous section, with only one of the top scoring gene drive designs (out of the 21 considered) producing viable heterozgyotic larvae (see Figure 7).

**Figure 7:**
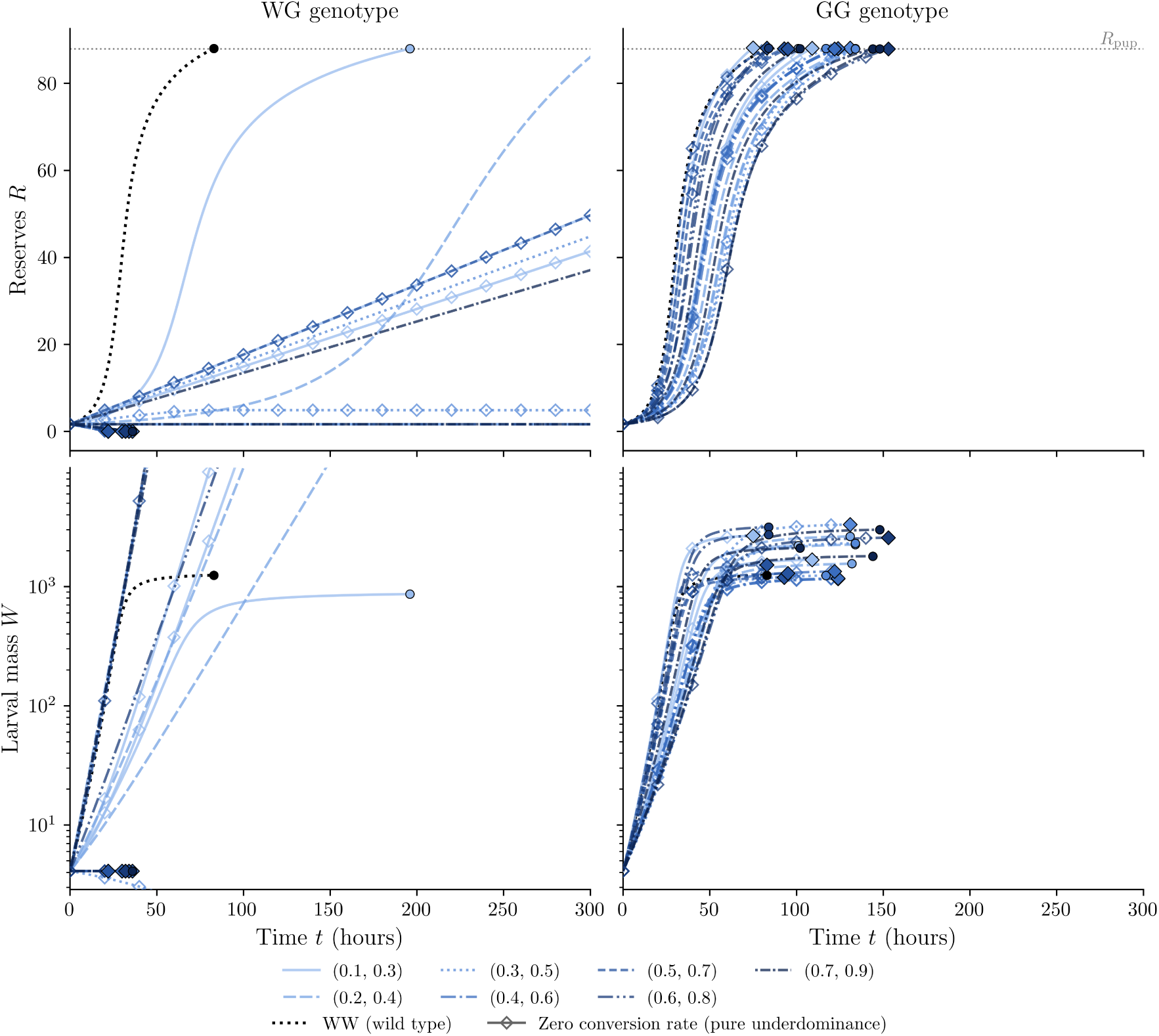
Mass (*W*) and reserves (*R*) trajectories over time for heterozygous (WW) and homozygous (GG) larvae, assuming unlimited food access, coloured by release threshold (the top three highest scoring gene drives are shown for each release threshold). Wild-type dynamics (black dotted line) are shown as a comparison, and gene drive designs corresponding to gene drives with zero conversion rate are highlighted. Trajectories are plotted until starvation *R* = 0, pupation *R* = *R*_pup_, or the end of the cycle, whichever comes first.

Furthermore, these results highlight that gene drive designs boost the fitness of GG organisms by reallocating food intake to increasing weight and size over accumulating reserves, resulting in greater competitive success at later times. As these gene drive designs use the same approach to boost the fitness of GG organisms, this suggests that successful gene drive designs will have similar ecological/physiological effects across target critical thresholds, but will also employ non-zero conversion rates for target critical thresholds away from 0.5.

### 3.4 Approximation of the Agent-Based Model

With the gene drive design with the highest average score across both objectives, we trained a deterministic surrogate model to approximate the dynamics of this gene drive and compared the results to a held-out test set of 2000 simulations. To confirm that our deterministic surrogate accurately approximates the dynamics of the agent-based model, we computed the root-mean-square error. We found that the deterministic surrogate has a root-mean-square-error of approximately 13.6, or less than 5% of the observed variation in WW or GG, as well as qualitatively reproducing the observed behaviour of the agent-based model (see Figure 8).

**Figure 8:**
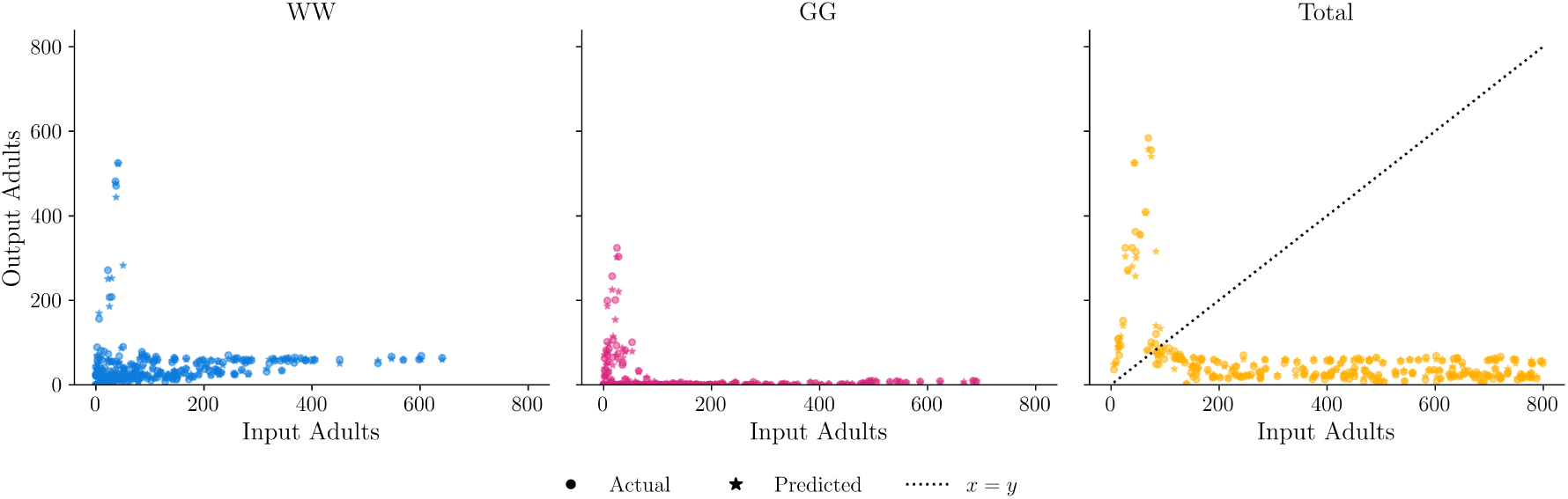
Comparison of the agent-based model and the neural network approximation for a randomly generated test set (only every 5th point is plotted to increase the readability of the figure). Each point represents the outcome of a simulation where a randomly generated population of adults are allowed to mate and lay eggs into the environment (the number of adults of each genotype is shown on the x-axis) and the number of adults of each genotype who survive to adulthood is recorded (shown on the y-axis). Note that survival is highly density-dependent, with low initial populations (less than 100 adults) leading to large population increases, but that survival falls sharply as populations sizes increase and the total number of larvae surviving to adulthood is fixed at approximately 100 for large initial populations (the line *x* = *y*, representing a fixed survival rate, is plotted as a comparison). Since the gene drive design approximated here has non-viable heterozygotes, the amount of WG organisms outputted was less than one for both the original model and the deterministic surrogate, so these have been omitted from the figure to improve readability.

For the stochastic neural spline flow (NSF) surrogate model, we validated the surrogate model by comparing the distribution of the test set outputs (e.g. the number of WW, WG, and GG individuals conditional on the known inputs) to the distribution of outputs generated by our NSF, using the inverse function approach developed by Linhart et al. [20]. We observed that the distribution was well calibrated for WW, but that the distribution of the GG population in the test set was slightly under-dispersed (it has a more sharply peaked distribution than the expected standard normal, implying that it is less variable than samples from the surrogate model). This suggests that the stochastic surrogate model may overstate the variability in the distribution of GG individuals (see Figure 9).

**Figure 9:**
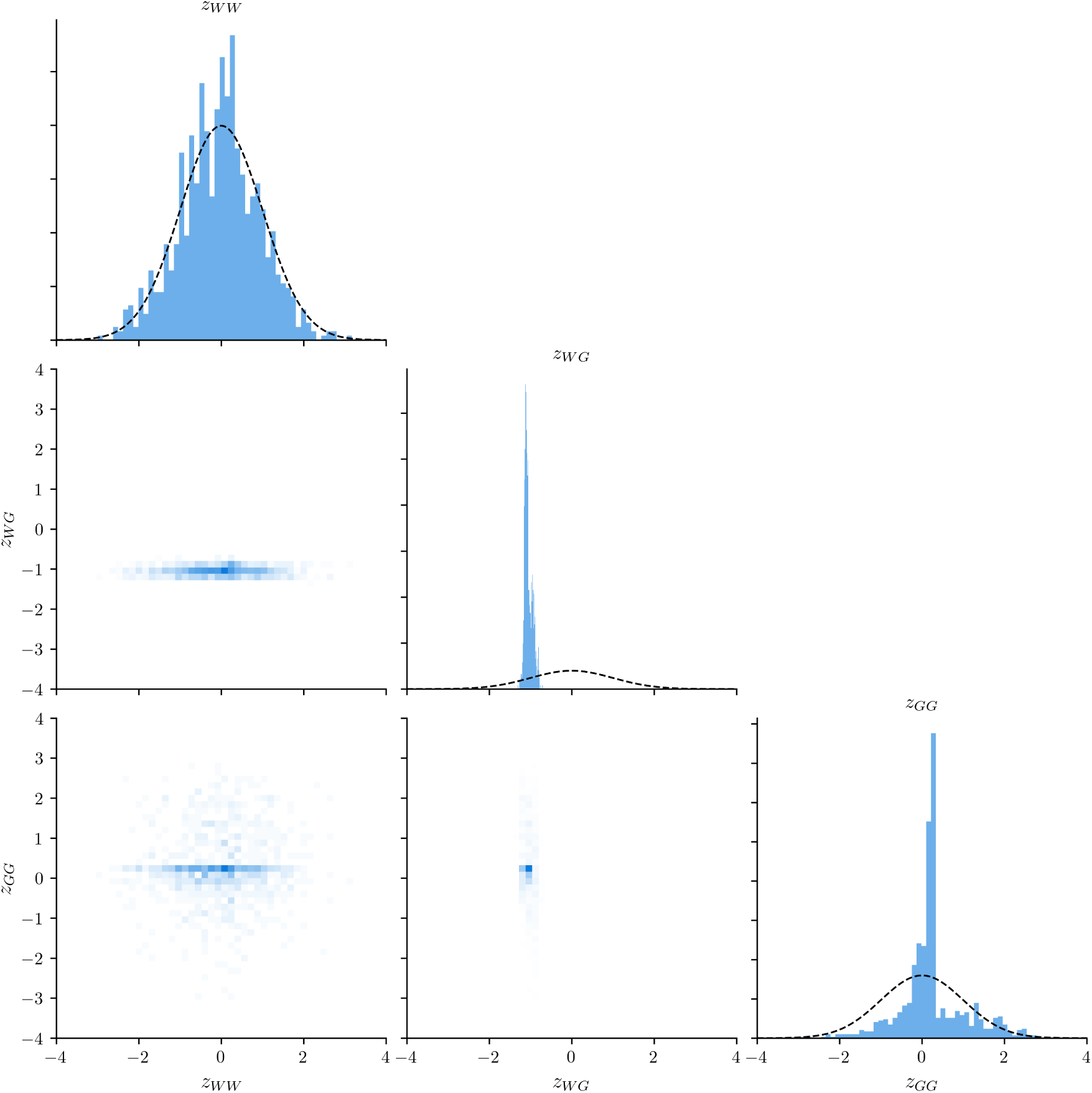
The latent distribution of the test set, with a standard normal plotted for reference. Note that the degenerate distribution of WG is expected, as the outputted number of heterozygotes is everywhere zero in our test and train datasets (this is because the gene drive design we are approximating has non-viable heterozygotes).

However, while these validation checks show generally good agreement between our surrogate models and the test dataset, they show only the accuracy of the model for a single generation (e.g. the ability to predict the number of adults at the next generation given the current number). To understand whether accumulated error in predictions led to inaccurate results in a multi-generation setting, we simulated populations over 10 generations from an initial starting point of 50 WW and 50 GG adults. To account for the stochastic nature of the agent-based model, we computed 50 trajectories for this model and report both the individual trajectories and the average over time. We expected that, given that this initial condition is near the target critical threshold, in some trajectories the gene drive will fix in the population and in some trajectories the gene drive will become extinct. Consequently, we expect that the feed-forward architecture may struggle to predict how the population evolves from this initial starting point, as the moment closure approximation (Equation 7) is less likely to be accurate in these settings where the variance of outcomes is high. In fact, while the feed-forward architecture performs well in the single generation test conducted above, it did not accurately capture the dynamics we observed in multi-generation simulations (see Figure 10).

**Figure 10:**
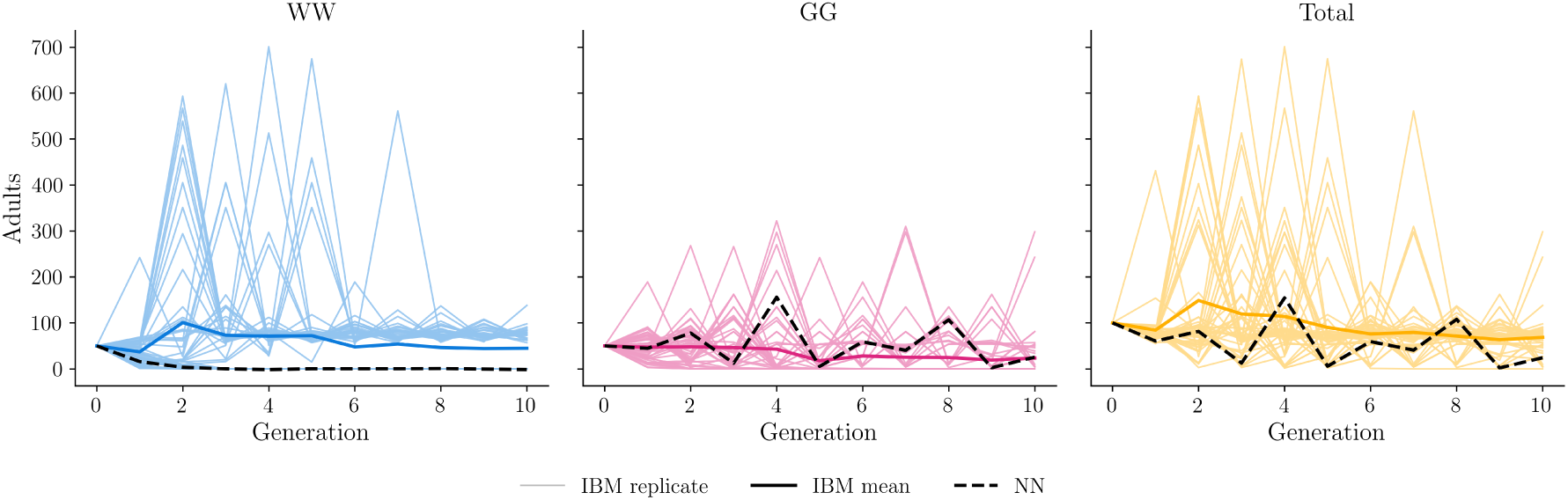
Comparison of the agent-based model and the neural network approximation over 10 generations, showing the abundance of adults of each genotype (and the total population) in each generation. Fifty replicates of the individual-based model are shown (thin coloured lines), with the mean plotted as a thick coloured line. Predictions from the neural network (e.g. the deterministic surrogate) are shown as a thick black dotted line. Note that the Neural Network predicts the subset of solutions where the gene drive fixes in the population, but does not capture the mean solution accurately. As in Figure 8 heterozygotes (genotype WG) are non-viable for this gene drive, and are consequently not plotted.

Considering now the NSF approximation, we found that the mean trajectory obtained by averaging over fifty replicates of the NSF was qualitatively similar to that obtained by averaging over fifty replicates from the agent-based model. Furthermore, the set of NSF replicates included trajectories where the gene drive became extinct and trajectories where the gene drive fixed in the population (see Figure 11), suggesting that this model could reproduce both possible outcomes observed in the agent-based model.

**Figure 11:**
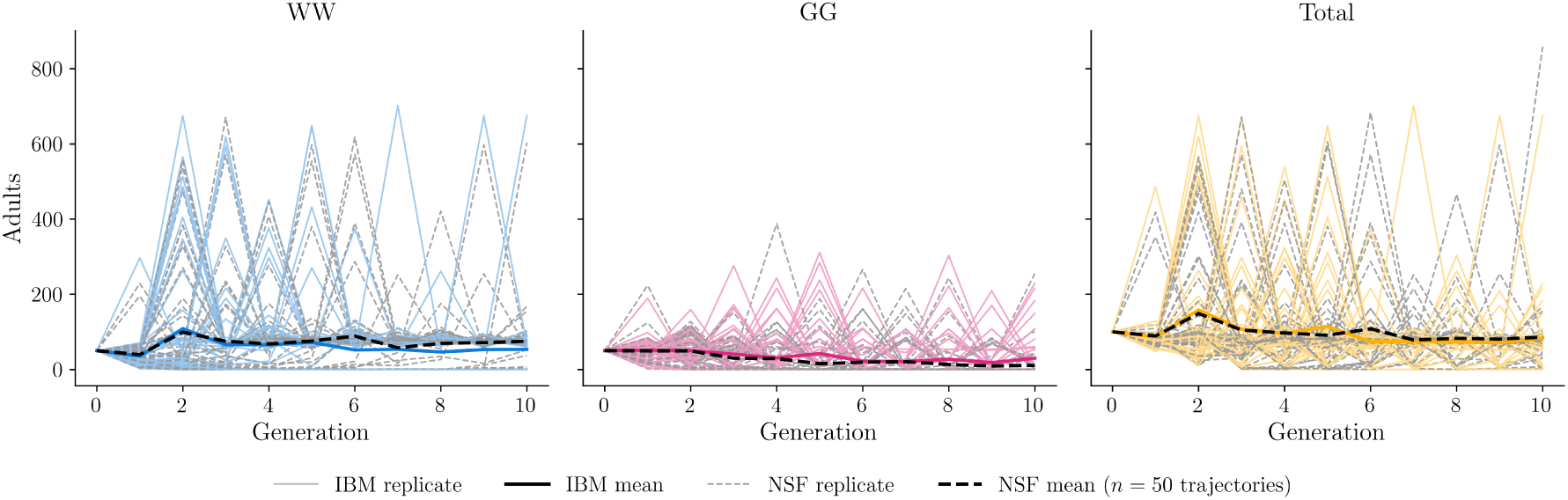
Comparison of the agent-based model and the NSF approximation over 10 generations, showing the abundance of adults of each genotype (and the total population) in each generation. Fifty replicates of the individual-based model are shown (thin coloured lines), with the mean plotted as a thick coloured line. Individual replicates from the NSF (e.g. the stochastic surrogate) are shown as thin black dotted lines, with the mean plotted as a thick black dotted line. Note that the approximation now captures both types of trajectories well. As in Figure 8 and Figure 10 heterozygotes (genotype WG) are non-viable for this gene drive, and are consequently not plotted.

These results demonstrate that both deterministic and stochastic surrogate models can effectively capture the dynamics of the agent-based model, but that only stochastic surrogate models are suitable for multi-generation simulations.

## 4 Discussion and Future Work

In this study we have investigated how ecological factors determine the performance of threshold dependent drives. Our results above highlight the importance of ecology in determining gene drive performance and the importance of building models that capture the interactions between genetics and ecology, as suggested by prior work [18, 35]. We find that ecological parameters governing the behaviour of the larvae have a dominant role in determining gene drive performance across much of the parameter space, but that genetic parameters have an important role in determining gene drive performance at the frontier of the best designs possible (the Pareto frontier). Our results highlight the importance of understanding the fitness costs of candidate gene drives in ecologically plausible settings, although these measurements are not always conducted in practice [21] and the best way to assess these costs is not yet known [17].

On this Pareto frontier, following [9] we find that efficient gene drive designs produce non-viable heterozygotes. In the case where a critical threshold of 0.4 is targeted (as suggested by [31]) we find that optimal gene drive designs use a variety of techniques to achieve success. Gene drive designs included both homing and non-homing designs, with the highest scoring gene drive design incorporating no homing component. We observe that these drives achieve competitive advantage by reallocating food to weight gain to increase the feeding success of GG organisms at late times. Changing release thresholds modified the optimal parameter set but the general design of gene drives remained similar except that non-zero conversion rates (e.g. homing gene drive components) were much more common for high and low release thresholds.

Our results suggest that developing detailed agent-based models of larval behaviour can be a useful tool to understand how different gene drive designs will behave. These detailed agent-based models can explore how gene drive designs may interact with density-dependent competition and serve as a bridge between population-scale models like MGDrivE 2.0 [34] or Skeeter Buster [22] and experimental trials. While our agent-based model is computationally costly compared to population-scale models, we find that stochastic surrogate models are an effective substitute which can capture the variation in outcomes observed across initial conditions. However we find that deterministic surrogate models are less effective in multi-generation settings, as such models depend on a moment-closure assumption that breaks down when there is significant variability in outcomes (such as near the critical threshold, where the gene drive is similarly likely to become extinct or fix in the population).

There are a number of ways in which this work could be extended to more accurately capture mosquito dynamics. Firstly, in the current model, only a single breeding site is considered. While this matches the laboratory setting in which gene drive research is often conducted, future work which extends this model to a setting with multiple breeding sites would be valuable, as spatial dynamics likely play an important role in determining gene drive success in ecologically plausible contexts [6].

Furthermore, our model assumes non-overlapping generations and that adult mosquitoes mate only once. While this assumption significantly simplifies model design, a more detailed framework with overlapping generations and the inclusion of more realistic mosquito mating behaviour would improve the verisimilitude of the model and allow for exploration of ecological/behavioural factors that are currently ignored by the model. For example, developing this approach would enable modelling to account for selection against WG or GG organisms at the time of mating, which has previously been demonstrated during field trials of genetically modified mosquitoes [13]. We expect that a model with overlapping generations and a more detailed treatment of mosquito mating behaviour (potentially including selection against WG or GG organisms) would still produce similar gene drive designs and highlight the importance of ecological parameters in determining results, but that negative selection during mating would generally increase the critical threshold for spread associated with a given gene drive design.

## 5 Conclusion

In this study, we modified an agent-based model of larval behaviour to incorporate a simple genetic framework. Using Bayesian optimisation, we optimised over the possible space of per-parameter genetic penalties to find optimal gene drives given a desired critical threshold for fixation. Our results provide insight into the mechanisms by which gene drive designs produce threshold dependent dynamics, suggesting that ecological and physiological parameters dominate over genetic parameters for much of the space. We further utilised machine learning techniques to develop stochastic and deterministic surrogate models for the agent-based model, allowing for rapid assessment of different scenarios. A promising route forward for ecologically-grounded gene drive modelling could combine agent-based models designed to capture behaviour that occurs at small scales (such as larval competition) with machine learning techniques to produce efficient approximate models. These efficient and ecologically grounded models of small-scale processes could then be used as building blocks for models that operate at the scale of large epidemiologically relevant landscapes.

## Acknowledgements

We thank Ziqian Xu, Professor Robin Thompson, and Professor Owen Lewis for insightful discussions and feedback on draft versions of this article.

## A Data and Code Availability

All data and code for this paper are available on Zenodo: https://doi.org/10.5281/zenodo.23914062

## B Environmental/Fixed Parameters for the Individual Based Model

The environmental and fixed parameters for the agent-based model govern a variety of aspects of the model, including the current temperature, the amount of resources available (and the rate at which these resources are replenished), the maximum number of eggs that can be laid at the breeding site, and number of larvae that can feed effectively at a given location. We chose typical parameter values from [36] which were based on empirical data from the literature. These parameters are summarised in Table 2.

**Table 2:** Default environmental and structural parameters for the agent-based model, reported in the model’s internal units. The larval-behaviour parameters (*a*, *α*, *β*, *c*, *g*) are given separately in Table 1.

| Symbol | Description | Default value |
|---|---|---|
| *Spatial structure* |  |  |
| — | Side length of the square breeding-site grid (cells) | 20 |
| *V*<sub>cell</sub> | Volume associated with each grid cell | 5 |
| — | Larval density level used to set the egg carrying capacity | 1.4 |
| — | Pond egg carrying capacity (*V*<sub>cell</sub> · density · 20<sup>2</sup>) | 2800 |
| *Temporal resolution and temperature* |  |  |
| — | Number of (hourly) time steps per cycle | 300 |
| *T* | Temperature | 20 °C |
| *Food and feeding* |  |  |
| — | Feeding mode | interference |
| *n*<sub>winners</sub> | Maximum number of larvae that may feed per cell | 4 |
| — | Initial food per cell (2700 × *n*<sub>winners</sub>) | 10800 |
| — | Food refill per cell every 24 steps (75 × *n*<sub>winners</sub>) | 300 |
| — | Assimilation efficiency (fraction of food assimilated) | 0.25 |
| *Larval physiology and life history* |  |  |
| *W*<sub>0</sub> | Initial larval weight | 4.1 |
| *R*<sub>0</sub> | Initial larval reserve (0.4 *W*<sub>0</sub>) | 1.64 |
| *R*<sub>th</sub> | Reserve set-point in the feeding response | 15 |
| *σ* | Width of the Gaussian feeding response in reserves | 23 |
| *R*<sub>pup</sub> | Reserve threshold for commitment to pupation | 87.9 |
| — | Delay between pupation commitment and pupation (time steps) | 37.5 |
| — | Egg to larva survival probability (for 20 ≤ *T* ≤ 30) | 0.9 |
| *Reproduction* |  |  |
| — | Eggs laid per female per cycle | 30 |

## C Implementation Details for Bayesian Optimisation

To implement the multi-objective Bayesian optimisation, we used the BoTorch library [2]. We defined the two objective functions as described in subsection 2.2, and used a Gaussian process surrogate model with a Matern kernel to model the relationship between the gene drive parameters and the objective function values. We used the q-Noisy Expected Hypervolume Improvement acquisition function to select the next set of gene drive parameters to evaluate at each iteration of the optimisation. We ran the optimisation for 150 iterations, starting from an initial set of 128 randomly sampled points in the parameter space. To control for noise in the stochastic simulations of the agent-based model, we evaluated each set of gene drive parameters on five replicates and used the mean outcomes as the objective function values for the optimisation. We batched the evaluation of the agent-based model to speed up the optimisation process, evaluating *q* = 4 sets of gene drive parameters in parallel at each iteration. To optimise the acquisition function we used the default LBFGS-B algorithm implemented in BoTorch, with 40 restarts and 2048 raw samples. Further details of the implementation can be found in the code available at https://doi.org/10.5281/zenodo.23914062.

## D Implementation Details for Neural Network Approximation

We used a feedforward neural network with 4 hidden layers and 1024 neurons per layer. To train the neural network, we generated a dataset of 40,000 samples of the input-output pairs of the agent-based model, where the inputs were the number of adults of each genotype at generation *t* and the outputs were the number of larvae of each genotype that survive to adulthood at generation *t* + 1. We trained the neural network on this dataset using the Adam optimizer with a learning rate of 0.0001 and a batch size of 128 for 900 epochs, reducing the learning rate by a factor of 10 every 300 epochs. We used mean squared error as the loss function. After training, we evaluated the performance of the neural network on a held-out test set of 2000 samples, finding that it provided a good approximation to the agent-based model (see Figure 8). Further details of the implementation can be found in the code available at https://doi.org/10.5281/zenodo.23914062.

## E Implementation Details for Neural Spline Flow Approximation

We used a neural spline flow (NSF, for details see [12]) to approximate the full distribution of the number of larvae of each genotype that survive to adulthood given an initial population of adults laying eggs into a breeding site. We used the Zuko package [27] to implement the NSF, using 5 hidden layers and 1024 features per layer. We trained the NSF using the Adam optimizer with a learning rate of 0.0005 and a batch size of 128 for 300 epochs, using the same test and train data as in Appendix D. We used the negative log likelihood as the loss function. After training, we evaluated the performance of the NSF on a held-out test set, finding that it provided a good approximation to the distribution of outcomes of the agent-based model. Further details of the implementation can be found in the code available at https://doi.org/10.5281/zenodo.23914062.

## F Construction of the Inverse Flow

To understand the construction of *π̃^−^*^1^, it is necessary to understand how neural spline flows function in more detail. A brief introduction is given here, but interested readers should consult [12] and [20] for more details. To generate an approximate distribution *π̃* (potentially conditioned on some vector of variables **y**), the NSF trains a neural network to construct an invertible mapping *φ*(**z**, **y**) such that:

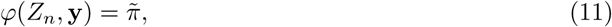

where *Z_n_* a standard normal distribution with the same dimension as *π̃*. Since this mapping is by construction invertible, it is then straightforward to construct *π̃^−^*^1^ = *φ^−^*^1^ which will by definition fufill the requirements given in Equation 10.

## G 2nd Order Sobol’ Indices

**Figure 12:**
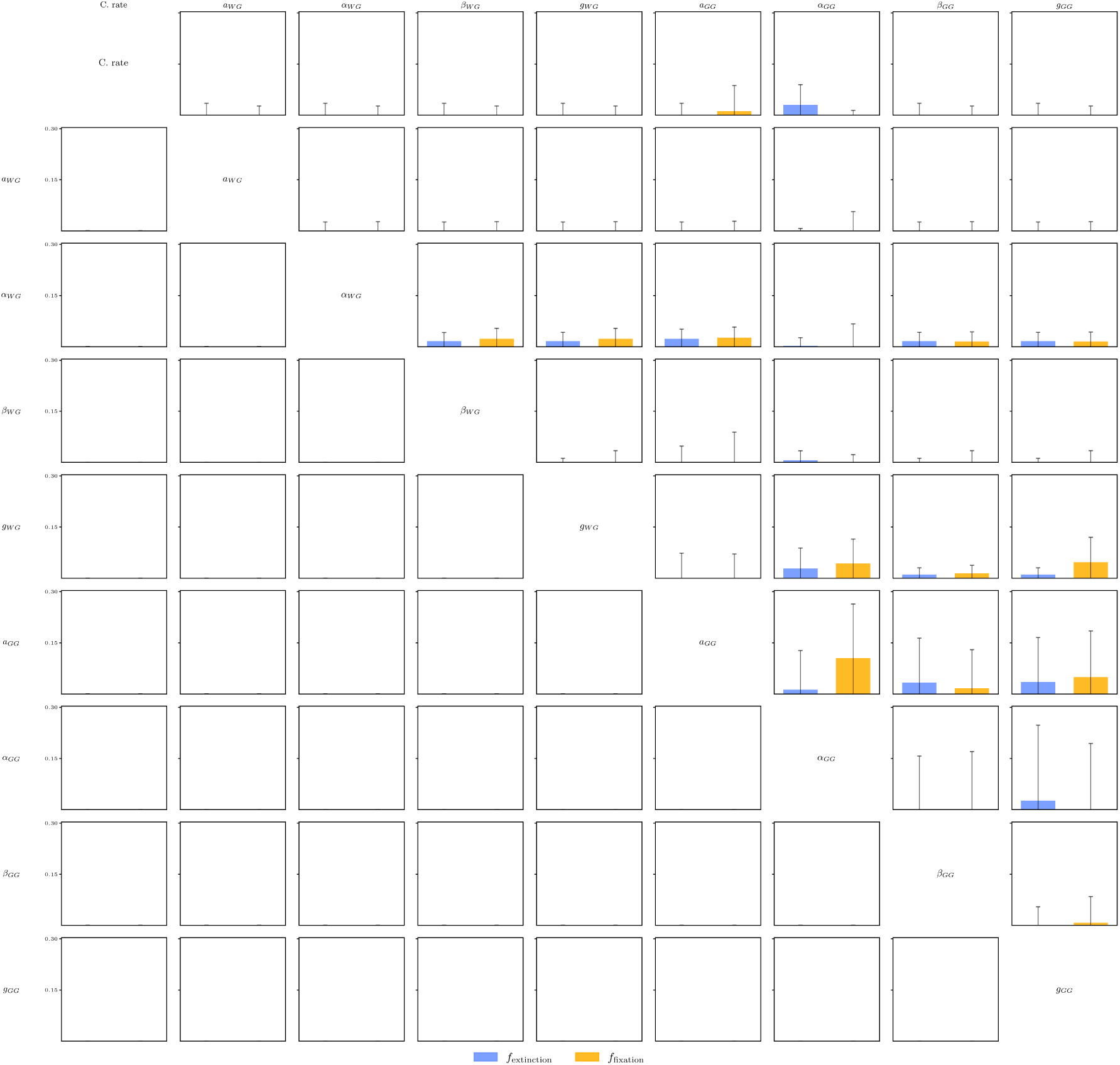
2nd order Sobol’ indices for each pair of parameters and objective function, calculated. Note that Sobol’ indices for most interactions are close to zero, except those between parameters determining the fitness of organisms homozygous for the gene drive.

## References

[1] Luke Alphey, Andrew McKemey, Derric Nimmo, Marco Neira Oviedo, Renaud Lacroix, Kelly Matzen, and Camilla Beech. Genetic control of Aedes mosquitoes. Pathog. Glob. Health, 107 (4):170–179, 2013. URL 10.1179/2047773213Y.0000000095.

[2] Maximilian Balandat, Brian Karrer, Daniel R. Jiang, Samuel Daulton, Benjamin Letham, Andrew G Wilson, and Eytan Bakshy. BoTorch: a framework for efficient Monte-Carlo Bayesian optimization. Adv. Neural Inf. Process. Syst., 33, 2020.

[3] Nick H. Barton and Michael Turelli. Spatial waves of advance with bistable dynamics: cytoplasmic and genetic analogues of Allee effects. Am. Nat., 178(3):E48–E75, 2011. URL 10.1086/661246.

[4] Samir Bhatt, Peter W. Gething, Oliver J. Brady, Jane P. Messina, Andrew W. Farlow, Catherine L. Moyes, John M. Drake, John S. Brownstein, Anne G. Hoen, Osman Sankoh, et al. The global distribution and burden of dengue. Nature, 496:504–507, 2013. URL 10.1038/nature12060.

[5] Marcia C. Castro, Mary E. Wilson, and David E. Bloom. Disease and economic burdens of dengue. Lancet Infect. Dis., 17(3):E70–E78, 2017. URL 10.1016/s1473-3099(16)30545-x.

[6] Jackson Champer, Joanna Zhao, Samuel E. Champer, Jingxian Liu, and Philipp W. Messer. Population dynamics of underdominance gene drive systems in continuous space. ACS Synth. Biol., 9(4):779–792, 2020. URL 10.1021/acssynbio.9b00452.

[7] Jackson Champer, Samuel E. Champer, Isabel K. Kim, Andrew G. Clark, and Philipp W. Messer. Design and analysis of CRISPR-based underdominance toxin-antidote gene drives. Evo. App., 14(4):1052–1069, 2021. URL 10.1111/eva.13180.

[8] Samuel Daulton, Maxmimilian Balandat, and Eytan Bakshy. Parallel Bayesian optimization of multiple noisy objectives with expected hypervolume improvement. Adv. Neural Inf. Process. Syst., 34, 2021.

[9] Stephen Davis, Nicholas Bax, and Peter Grewe. Engineered underdominance allows efficient and economical introgression of traits into pest populations. J. Theor. Biol., 212(1):83–98, 2001. URL 10.1006/jtbi.2001.2357.

[10] Jean-Marc O. Depinay, Charles M. Mbogo, Gerry Killeen, Bart Knols, John Beier, John Carlson, Jonathan Dushoff, Peter Billingsley, Henry Mwambi, John Githure, et al. A simulation model of African Anopheles ecology and population dynamics for the analysis of malaria transmission. Malar. J., 3(29), 2004. URL 10.1186/1475-2875-3-29.

[11] Sumit Dhole, Alun L. Lloyd, and Fred Gould. Gene drive dynamics in natural populations: The importance of density dependence, space, and sex. Annu. Rev. Ecol. Evol. Syst., 51:505–531, 2020. URL 10.1146/annurev-ecolsys-031120-101013.

[12] Conor Durkan, Artur Bekasov, Iain Murray, and George Papamakarios. Neural spline flows. Adv. Neural Inf. Process. Syst., 32, 2019.

[13] Luca Facchinelli, Laura Valerio, Janine M. Ramsey, Fred Gould, Rachael K. Walsh, Guillermo Bond, Michael A. Robert, Alun L. Lloyd, Anthony A. James, Luke Alphey, et al. Field cage studies and progressive evaluation of genetically-engineered mosquitoes. PLoS Negl. Trop. Dis., 7(1):e2001, 2013. URL 10.1371/journal.pntd.0002001.

[14] Penelope A. Hancock, Vanessa L. White, Scott A. Ritchie, Ary A. Hoffmann, and H. Charles J. Godfray. Predicting Wolbachia invasion dynamics in Aedes aegypti populations using models of density-dependent demographic traits. BMC Biol., 14:96, 2016. URL 10.1186/s12915-016-0319-5.

[15] Penelope A. Hancock, Scott A. Ritchie, Constantianus J.M. Koenraadt, Thomas W. Scott, Ary A. Hoffmann, and H. Charles J. Godfray. Predicting the spatial dynamics of Wolbachia infections in Aedes aegypti arbovirus vector populations in heterogeneous landscapes. J. Appl. Ecol., 56(7):1674–1686, 2019. URL 10.1111/1365-2664.13423.

[16] Trinh Manh Hung, Donald S. Shepard, Alison A. Bettis, Huyen Anh Nguyen, Angela McBride, Hannah E. Clapham, and Hugo C. Turner. Productivity costs from a dengue episode in Asia: a systematic literature review. BMC Infect. Dis., 20:393, 2020. URL 10.1186/s12879-020-05109-0.

[17] Anna Janzen, Ratnasri Pothula, Adam Sychla, Nathan R. Feltman, and Michael J. Smanski. Predicting thresholds for population replacement gene drives. BMC Biol., 22:40, 2024. URL 10.1186/s12915-024-01823-2.

[18] Jaehee Kim, Keith D. Harris, Isabel K. Kim, Shahar Shemesh, Philipp W. Messer, and Gili Greenbaum. Incorporating ecology into gene drive modelling. Ecol. Lett., 26(S1):S62–S80, 2023. URL 10.1111/ele.14194.

[19] Philip T. Leftwich, Matthew P. Edgington, Tim Harvey-Samuel, Leonela Z. Carabajal Paladino, Victoria C. Norman, and Luke Alphey. Recent advances in threshold-dependent gene drives for mosquitoes. Biochem. Soc. Trans., 46(5):1203–1212, 2018. URL 10.1042/BST20180076.

[20] Julia Linhart, Alexandre Gramfort, and Pedro Rodrigues. L-c2st: Local diagnostics for posterior approximations in simulation-based inference. Adv. Neural Inf. Process. Syst., 36, 2023. URL 10.52202/075280-2460.

[21] Vanessa M. Macias, Johanna R. Ohm, and Jason L. Rasgon. Gene drive for mosquito control: where did it come from and where are we headed? Int. J. Env. Res Public Health, 14(9):1006, 2017. URL 10.3390/ijerph14091006.

[22] Krisztian Magori, Mathieu Legros, Molly E. Puente, Dana A. Focks, Thomas W. Scott, Alun L. Lloyd, and Fred Gould. Skeeter Buster: a stochastic, spatially explicit modeling tool for studying Aedes aegypti population replacement and population suppression strategies. PLoS Negl. Trop. Dis., 3(9):e508, 2009. URL 10.1371/journal.pntd.0000508.

[23] Ace R. North, Austin Burt, and H. Charles J. Godfray. Modelling the potential of genetic control of malaria mosquitoes at national scale. BMC Biol., 17:26, 2019. URL 10.1186/s12915-019-0645-5.

[24] Marcelo Otero, Nicolás Schweigmann, and Hernán G Solari. A stochastic spatial dynamical model for Aedes aegypti. Bull. Math. Biol., 70:1297–1325, 2008. URL 10.1007/s11538-008-9300-y.

[25] Robyn Raban, Anthony A. James, and Omar S. Akbari. Advances in CRISPR gene drives for mosquito population control. Curr. Opin. Microbiol., 90:102712, 2026. URL 10.1016/j.mib.2026.102712.

[26] Gordana Rašić and John M. Marshall. Integrating mosquito genomics into simulation modeling: opportunities for better-informed biocontrol. Curr. Opin. Insect Sci., 73:101456, 2026. URL 10.1016/j.cois.2025.101456.

[27] François Rozet et al. Zuko: Normalizing flows in pytorch. Software Package, 2022. URL https://pypi.org/project/zuko.

[28] Frederic W. Selck, Amesh A. Adalja, and Crystal R. Boddie. An estimate of the global health care and lost productivity costs of dengue. Vector Borne Zoonotic Dis., 14(11):824–826, 2014. URL 10.1089/vbz.2013.1528.

[29] Donald S. Shepard, Eduardo A. Undurraga, Yara A. Halasa, and Jeffrey D. Stanaway. The global economic burden of dengue: a systematic analysis. Lancet Infect. Dis., 16(8):935–941, 2016. URL 10.1016/s1473-3099(16)00146-8.

[30] Jeffrey D. Stanaway, Donald S. Shepard, Eduardo A. Undurraga, Yara A. Halasa, Luc E. Coffeng, Oliver J. Brady, Simon I. Hay, Neeraj Bedi, Isabela M. Bensenor, Carlos A. Castañeda-Orjuela, et al. The global burden of dengue: an analysis from the Global Burden of Disease Study 2013. Lancet Infect. Dis., 16(6):712–723, 2016. URL 10.1016/S1473-3099(16)00026-8.

[31] Hidenori Tanaka, Howard A. Stone, and David R. Nelson. Spatial gene drives and pushed genetic waves. Proc. Natl. Acad. Sci. U.S.A., 114(32):8452–8457, 2017. URL 10.1073/pnas.1705868114.

[32] Annelies Wilder-Smith, Thomas Cherian, and Joachim Hombach. Dengue vaccine development and deployment into routine immunization. Vaccines, 13(5):483, 2025. URL 10.3390/vaccines13050483.

[33] Adeline E. Williams, Alexander W.E. Franz, William R. Reid, and Ken E. Olson. Antiviral effectors and gene drive strategies for mosquito population suppression or replacement to mitigate arbovirus transmission by Aedes aegypti. Insects, 11(1):52, 2020. URL 10.3390/insects11010052.

[34] Sean L. Wu, Jared B. Bennett, Héctor M. Sánchez C., Andrew J. Dolgert, Tomás M. León, and John M. Marshall. MGDrivE 2: A simulation framework for gene drive systems incorporating seasonality and epidemiological dynamics. PLoS Comput. Biol., 17(5):e1009030, 2021. URL 10.1371/journal.pcbi.1009030.

[35] Ziqian Xu and Michael B. Bonsall. Ecology of gene drives: The role of density-dependent feedbacks on the efficacy and dynamics of two-locus underdominance gene drive systems. Evol. Appl., 18(3):e70079, 2025. URL 10.1111/eva.70079.

[36] Ziqian Xu and Michael B. Bonsall. Emergent density dependence: an individual-based model of larval competition in *Aedes aegypti* (Dipter: Culcidae). Under Review, 2026.

[37] Laith Yakob, Luke Alphey, and Michael B. Bonsall. Aedes aegypti control: the concomitant role of competition, space and transgenic technologies. J. Appl. Ecol., 45(4):1258–1265, 2008. URL 10.1111/j.1365-2664.2008.01498.x.

